# Transcriptome and epigenome diversity and plasticity of muscle stem cells following transplantation

**DOI:** 10.1101/2020.05.20.107219

**Authors:** Brendan Evano, Diljeet Gill, Irene Hernando-Herraez, Glenda Comai, Thomas M. Stubbs, Pierre-Henri Commere, Wolf Reik, Shahragim Tajbakhsh

## Abstract

Adult skeletal muscles are maintained during homeostasis and regenerated upon injury by muscle stem cells (MuSCs). A heterogeneity in self-renewal, differentiation and regeneration properties has been reported for MuSCs based on their anatomical location. Although MuSCs derived from extraocular muscles (EOM) have a higher regenerative capacity than those derived from limb muscles, the molecular determinants that govern these differences remain undefined. Here we show that EOM and limb MuSCs have distinct DNA methylation signatures associated with enhancers of location-specific genes, and that the EOM transcriptome is reprogrammed following transplantation into a limb muscle environment. Notably, EOM MuSCs expressed host-site specific positional *Hox* codes after engraftment and self-renewal within the host muscle. However, about 10% of EOM-specific genes showed engraftment-resistant expression, pointing to cell-intrinsic molecular determinants of the higher engraftment potential of EOM MuSCs. Our results underscore the molecular diversity of distinct MuSC populations and molecularly define their plasticity in response to microenvironmental cues. These findings provide insights into strategies designed to improve the functional capacity of MuSCs in the context of regenerative medicine.

## INTRODUCTION

Skeletal muscles are essential for physiological functions such as locomotion, breathing, and metabolism, and they represent up to 40% of the human body mass. Tissue-specific muscle stem cells (MuSCs) ensure skeletal muscle homeostasis and regeneration^1–3^. MuSCs have been implicated in the etiology of some muscular dystrophies^4,5^ as well as age-associated impaired muscle regeneration^6–11^ leading to an incapacitating decrease of muscle mass and strength^12–15^. Stem-cell therapies have proven to be challenging for muscular dystrophies, as they require delivering enough functional MuSCs to the right muscle groups and *ex vivo* amplification of healthy MuSCs results in a major decline in their regenerative capacity and stemness properties^16^.

Most of the knowledge about MuSC biology arises from the study of limb muscles, whereas MuSCs from other muscle groups remain less well characterized. Extraocular muscles (EOMs) are responsible for eye movements, with the basic pattern of 6 muscles conserved in most vertebrate species^17^. Limb muscles derive from the somitic mesoderm and rely in part on *Pax3* expression and function^18–20^. In contrast, EOMs derive from cranial mesoderm and rely initially on *Mesp1* and *Pitx2* for their emergence, yet unlike the majority of other cranial-derived muscles, their founder stem cell population^21,22^ arises independently of *Tbx1* function^17,23–29^. After the distinct specification of cranial and trunk progenitors, the core myogenic regulatory factors Myf5, Mrf4, Myod, and Myogenin regulate myogenic cell commitment and differentiation^30^. In adult homeostatic muscles, MuSCs are mostly quiescent. They are activated upon muscle injury, proliferate and differentiate to contribute to new muscle fibers, or self-renew to reconstitute the stem cell pool^31–34^. This process is accompanied by a temporal expression of cell fate markers, such as the transcription factors Pax7 (stem), Myod (commitment) and Myogenin (differentiation). Several reports indicate that EOMs are functionally different from their hindlimb counterparts since they are preferentially spared in ageing and several muscular dystrophies^35–38^. Interestingly, adult EOM-derived MuSCs cells have superior *in vitro* proliferation, self-renewal and differentiation capacities, as well as a superior *in vivo* engraftment potential, compared to limb-derived MuSCs^39^. These properties were maintained by EOM-derived MuSCs from dystrophic or aged mice^39^. However, the unique functional properties of adult EOM MuSCs remain undefined at the molecular level, as well as whether their specificity is instructed by cell-intrinsic factors or through interactions with their microenvironment (niche).

DNA methylation is a critical epigenetic mechanism involved in establishing and maintaining cellular identity. Dynamic changes in expression of DNA (de)methylation enzymes were reported in MuSCs during muscle regeneration^40–42^, indicating a potential increase of DNA methylation from quiescent to activated MuSCs. Several studies investigated the DNA methylation signatures of proliferating and differentiating cultures of MuSCs^43–45^ and reported some changes in DNA methylation patterns during late myogenic differentiation. Further, aged MuSCs show increased DNA methylation heterogeneity at promoters, associated with a degradation of coherent transcriptional networks^46^. However, whether quiescent MuSCs from homeostatic muscles at different anatomical locations display similar or different DNA methylation patterns remains unknown.

Here we performed parallel DNA methylation and transcriptome sequencing to characterize the specific identity of adult mouse EOM and limb MuSCs at the molecular level. We used heterotopic transplantation of EOM MuSCs into limb muscles to challenge their fate and assess their plasticity. We show that their specific identity is mostly niche-driven as they adopt a limb-like molecular signature. Nevertheless, we also identify EOM-specific genes that resist reprogramming by the microenvironment, indicating potential candidates to manipulate in limb-derived MuSCs for improving their regenerative capacity in the context of cellular therapies.

## RESULTS

To investigate the molecular differences between EOM and TA MuSCs *in vivo*, MuSCs from both locations were isolated by FACS (Fluorescence-Activated Cell Sorting) from 10 adult *Tg:Pax7-nGFP*^17^ mice and processed for RNA-sequencing and BS-sequencing as in^47,48^ (Figure 1A). Unsupervised Principal Component Analysis (PCA) on the transcriptomes revealed clear discrimination between TA and EOM MuSCs samples, indicating location-specific transcriptional identities (Figure 1B). To further investigate these transcriptional signatures, we performed differential expression analysis and found 261 genes significantly upregulated in EOM MuSCs (EOM Differentially Expressed Genes, DEGs) and 339 genes significantly upregulated in TA MuSCs (TA DEGs) (DE-seq, P <0.05, FC >2) (Figure 1C). The muscle stem cell marker *Pax7* and the myogenic factors *Myf5* and *Myod* showed slightly higher expression levels in TA compared to EOM MuSCs, while the transcription factor *Pitx2* showed higher expression levels in EOM MuSCs (Figure 1D). As reported^49^, *Pax3* was expressed in TA MuSCs and not detected in EOM MuSCs. Other genes specifically up-regulated in TA MuSCs included the developmental factors *Lbx1 and Tbx1.* Lbx1 is a homeobox transcription factor required for the migration of myogenic progenitor cells to the limbs, and Tbx1 is a T-box containing transcription factor necessary for the development of craniofacial muscles^50–52^. Examples of EOM up-regulated genes included the homeobox transcription factors *Lmx1a* and *Alx4*^30,53,54^ as well as *Mobp* (involved in myelination^55^) (Figure 1D). Further, genes of the *Hox* gene family were upregulated in TA samples (Figure 1E), consistent with their anteroposterior expression pattern in vertebrates^56^. Overall, 13 *Hox* genes were upregulated, representing 64% of the *HoxA* cluster and 75% of the *HoxC* cluster. Notably, these *Hox* genes are expressed more posteriorly along the body axis^57^. Overall, differentially expressed genes were enriched for developmental processes suggesting that location-specific patterns reflect their different origins during embryogenesis (Supplementary Figure 1).

**Figure 1.**
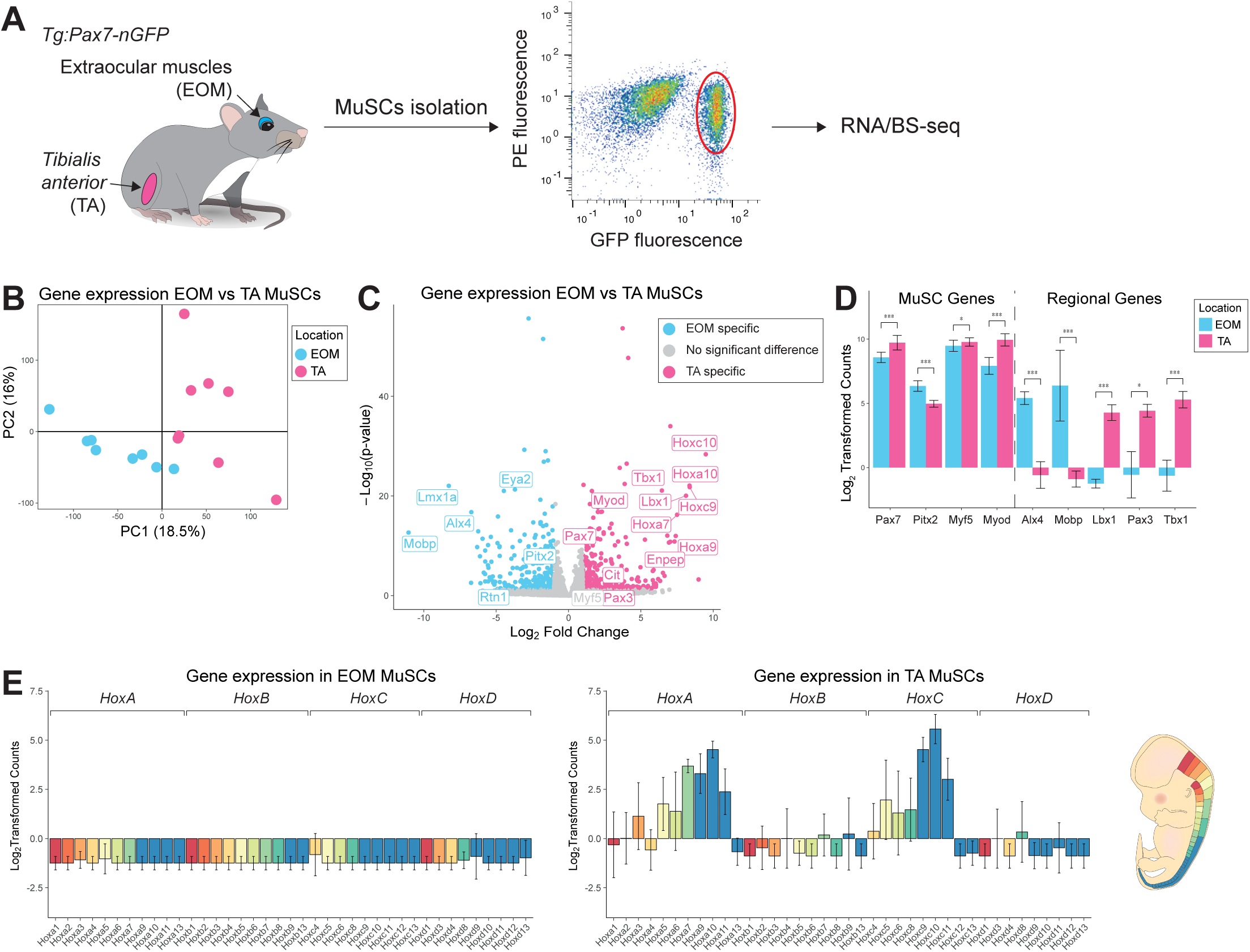
Head and limb-derived MuSCs display specific transcriptome signatures. (A) Experimental scheme. Head and limb MuSCs were isolated by FACS from EOM and TA muscles respectively from *Tg:Pax7-nGFP* adult mice and processed through RNA- and bisulfite-sequencing. N=10 mice, n=500 cells/mouse (Supplementary Table 1). (B) PCA analysis of EOM and TA MuSCs transcriptomes. The first principal component separates samples based on their anatomical location. (C) Volcano plot showing the results of differential expression analysis of EOM and TA MuSCs. Genes that demonstrated a fold change greater than 2 and a p-value less than 0.05 according to DESeq2 were classified as differentially expressed and colour coded by their tissue specificity. (D) Selected markers and differentially expressed genes between EOM and TA MuSCs. Error bars represent the standard deviation of the mean. p-values were determined by DESeq2. *** p < 0.001, ** p < 0.01, * p < 0.05. (E) Gene expression analysis of throughout the *HoxA, HoxB, HoxC* and *HoxD* clusters in EOM (left) and TA (middle) MuSCs. Genes were color-coded according to their antero-posterior expression domain in mouse at embryonic day 12.5 (right, adapted from^57^).

Overall, global DNA methylation levels were similar between EOM and TA MuSCs, with EOM MuSCs showing slightly higher levels of DNA methylation relative to TA MuSCs. On average genome-wide DNA methylation levels were around 50%, promoters and enhancers were hypomethylated (25% and 30% respectively) and repeat elements were hypermethylated (85%) (Figure 2A). We restricted our more detailed analysis to enhancers (regions marked by H3K27Ac in MuSCs that did not overlap promoters)^58^ and promoters (−2000bp to 500bp of the TSS of Ensembl genes) and calculated DNA methylation levels over each genomic element. PCA analysis for promoter regions showed no clear differences between TA and EOM MuSCs (Figure 2B). In contrast, a clear separation by anatomical location was observed when considering enhancer regions (Figure 2C). This clustering was also observed when restricting the analysis to 544 enhancers associated with TA and EOM DEGs (enhancer regions within 1 Mb to the closest transcription start site, Supplementary Figure 2A). Furthermore, enhancers associated with genes specifically upregulated in TA MuSCs were less methylated in TA MuSCs compared to EOM MuSCs. Similarly, enhancers associated with genes upregulated in EOM MuSCs were less methylated in EOM MuSCs compared to TA MuSCs (Figure 2D). To examine which enhancers were responsible for shifts in the distributions, we determined which enhancers were significantly differentially methylated using a rolling Z-score. Not all of the enhancers associated with TA or EOM DEGs were classified as differentially methylated, suggesting only a subset of these genes was regulated by enhancer methylation (Figure 2E). We also identified 29 Differentially Methylated Regions (DMRs) with more than 5 consecutive CpG methylation sites and 10% methylation differences. The majority of the DMRs overlapped enhancer or promoter regions such as the DMR overlapping the *Tbx1* promoter which is hypomethylated in TA MuSCs (Mean methylation levels in TA: 13%; mean methylation levels in EOM: 80%). These results suggest that DNA methylation patterns at regulatory regions contribute to the location-specific transcriptional profile of MuSCs.

**Figure 2.**
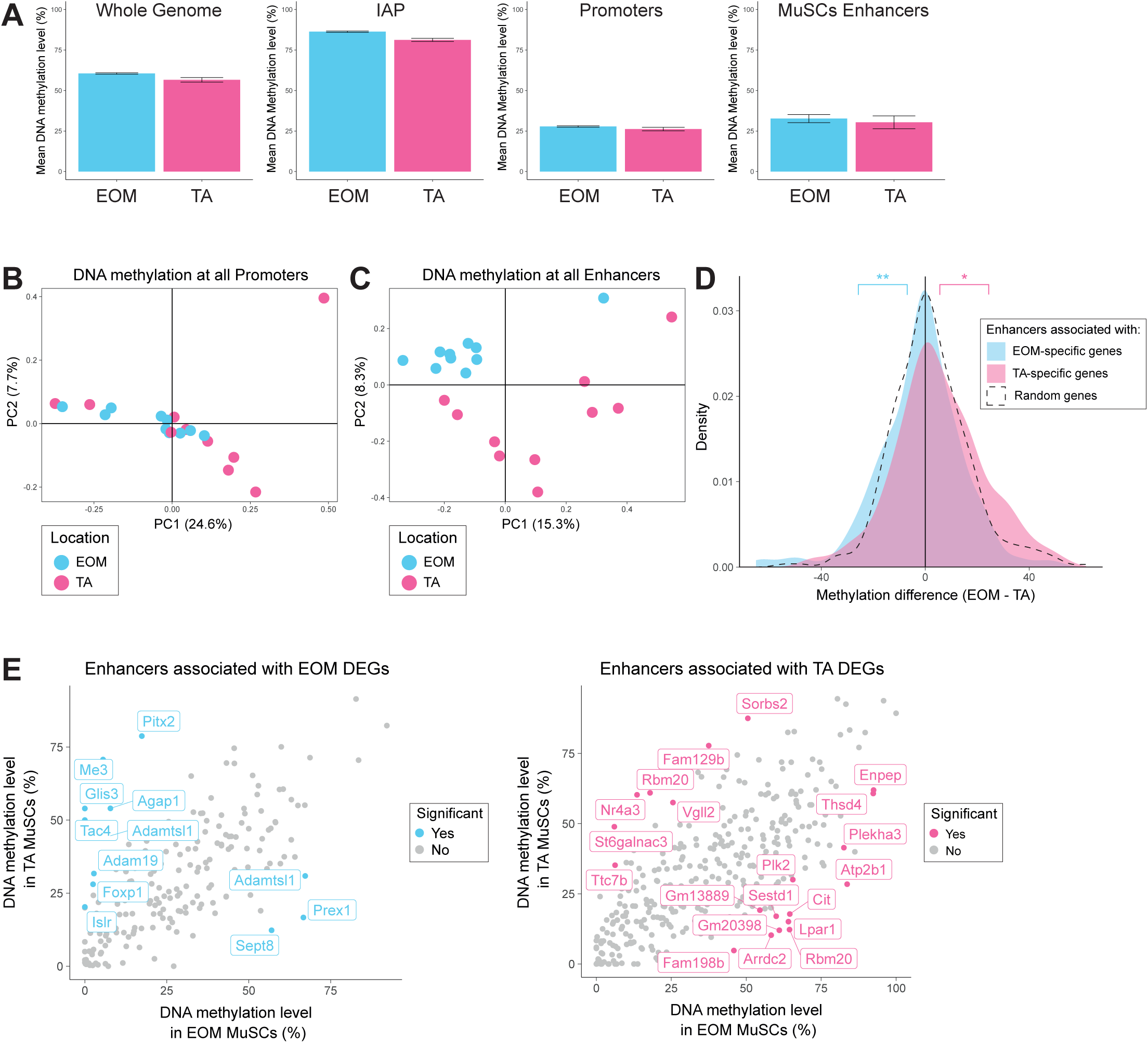
Head and limb-derived MuSCs display specific DNA methylation patterns at enhancers. (A) Mean DNA methylation levels of EOM and TA MuSCs across the whole genome, IAP repeat elements, promoters and enhancers. Promoters were defined as −2000bp to 500bp of the TSS of Ensembl genes. H3K27ac peaks were called using macs2 on H3K27ac ChIP-seq data obtained from^58^. Enhancers were defined as H3K27ac peaks that did not overlap promoters. Enhancers were linked to genes based on proximity. Overall DNA methylation was similar between the two locations, though EOM MuSCs had slightly higher levels of DNA methylation relative to TA MuSCs. (B) PCA analysis of DNA methylation at promoters fails to separate EOM and TA samples based on their anatomical location. (C) PCA analysis of DNA methylation at enhancers separates EOM and TA samples based on their anatomical location. (D) Density plots showing methylation differences between EOM and TA MuSCs at enhancers associated with location-specific genes (enhancers regions within 1 Mb to the closest transcription start site of a DEG). Enhancers associated with EOM genes (n=198) were significantly hypomethylated in EOM MuSCs relative to TA MuSCs when compared to a random subset of enhancers (n=350), while enhancers associated with TA genes (n=346) were significantly hypermethylated in the same comparison. ** p < 0.01, * p < 0.05 by Welch’s t test. (E) Scatter plots comparing the mean DNA methylation levels of enhancers associated with location DEGs in EOM and TA MuSCs. Enhancers were colour coded as significant if they were also found to be differentially methylated by a rolling Z score approach (p < 0.05) when comparing the methylation levels of all enhancers in EOM and TA MuSCs (Supplementary Figure 2B). The significant enhancers have been labelled with their associated gene. Only a subset of enhancers associated with location DEGs were differentially methylated. More enhancers associated with EOM DEGs were hypermethylated in TA MuSCs than in EOM MuSCs. Likewise, more enhancers associated with TA DEGs were hypermethylated in EOM MuSCs than in TA MuSCs.

To investigate MuSC plasticity and the influence of the cellular microenvironment on the observed location-specific signatures, we performed heterotopic transplantations of EOM MuSCs into TA muscles. Specifically, EOM MuSCs from *Tg:Pax7-nGFP* mice were transplanted into pre-injured TA muscles of immunodeficient *Rag2*^*-/-*^*;γC*^*-/-* 59^ mice. As a control, and for each donor mouse, TA MuSCs were transplanted into the same recipient mouse, in the contralateral pre-injured TA muscle. After 28 days when muscle regeneration is complete and self-renewal of MuSCs can be evaluated, engrafted EOM and TA MuSCs were re-isolated by FACS based on GFP positivity (post-graft samples). Additionally, a fraction of the EOM and TA MuSCs before grafting was kept as a control (pre-graft samples). Pre-graft and post-graft EOM and TA MuSCs were then processed for RNA-sequencing and BS-sequencing (Figure 3A). A total of 10 donor mice were analysed (Supplementary Table 1). We first observed important transcriptome differences between pre and post-graft MuSCs (Supplementary Figure 3A). Importantly, although muscle regeneration is generally considered to be largely achieved within 28 days post-injury^60^, transcriptome analysis showed substantial differences between pre-graft and post-graft TA MuSCs (Supplementary Figure 3B), indicating that our transplantation of MuSCs *per se* had a direct effect on gene expression. Genes upregulated after transplantation of TA MuSCs were enriched for immunological processes (Supplementary Figure 3C), including genes encoding interferon-induced proteins and complement proteins. However, as expected for this time point, the myogenic differentiation marker *Myogenin* was downregulated in the post-graft cells, indicating that cells at the time of re-isolation were not undergoing active myogenic commitment or differentiation. We also observed a slight global increase in the DNA methylation levels after transplantation (mean TA pre-graft: 56.6%, mean TA post-graft: 59.2%; Supplementary Figure 3D). Whether these observations reflect molecular changes with slow kinetics on MuSCs during muscle regeneration or are the result of the adaptation of engrafted MuSCs to an immunodeficient environment remains to be explored. The latter is favoured as recent RNA-seq analyses of MuSCs during regeneration showed that MuSCs re-acquire a quiescent homeostatic transcriptome as early as 7 days post-injury^61^.

**Figure 3.**
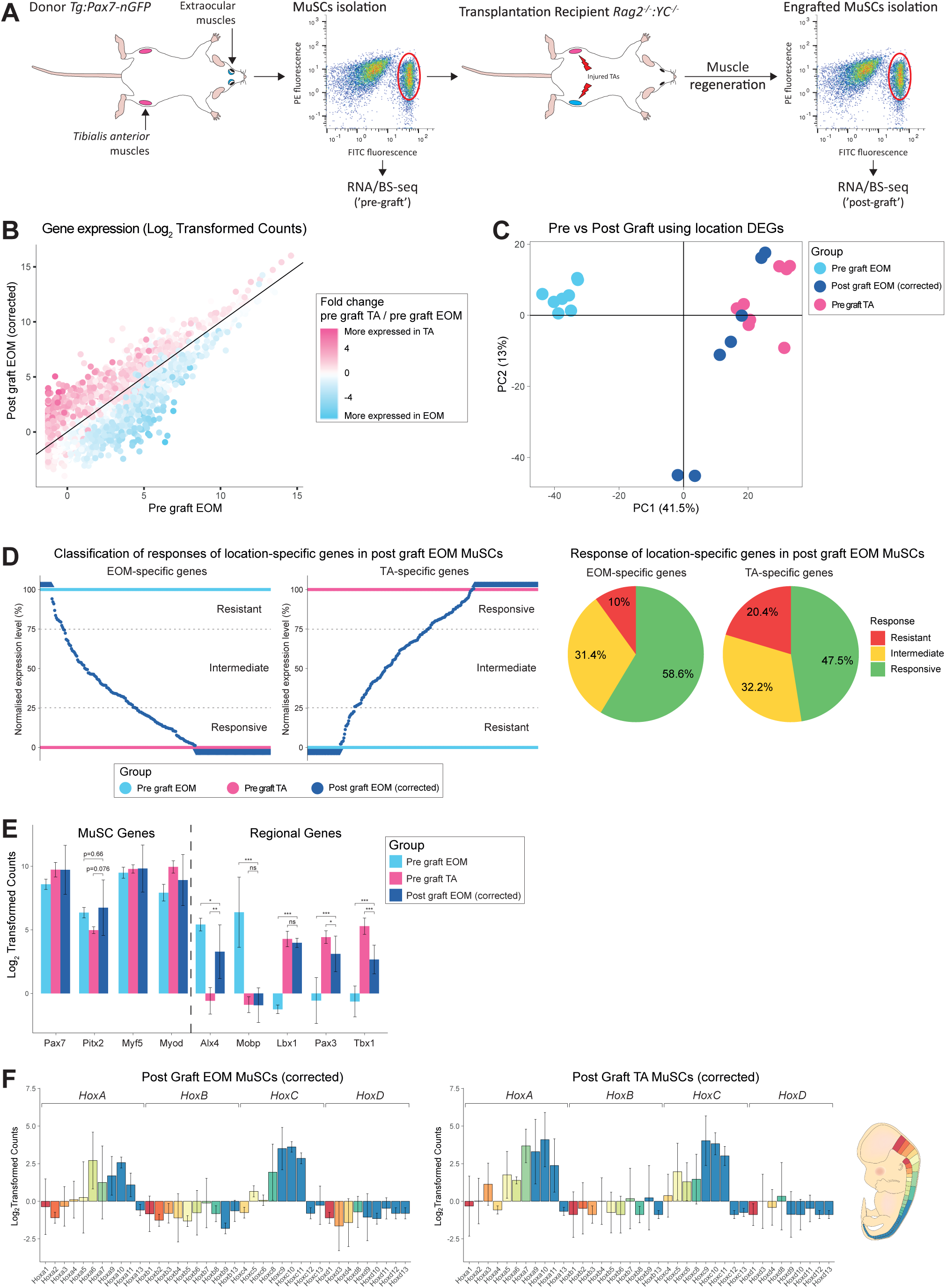
Head-derived MuSCs adopt a limb-like transcriptome upon transplantation in limb muscles. (A) Experimental scheme. EOM and TA-derived MuSCs were engrafted into pre-injured recipient TA muscles. After regeneration, engrafted MuSCs were re-isolated and processed through RNA- and BS-sequencing. ‘Post-graft’ MuSCs were compared to their ‘Pre-graft’ counterparts. N=10 donor mice and 10 recipient mice. For each donor mouse, equal number of EOM and TA MuSCs (in the range of 10,000 cells) were transplanted to TA muscles of the same recipient mouse. For each donor muscle, 500 pre-graft cells and 60 re-isolated post-graft cells (mean value) were analysed (Supplementary Table 1). (B) Expression analysis of EOM post-graft and EOM pre-graft. Each dot represents a gene. Genes are color-coded according to their fold change in pre-graft TA *vs* EOM MuSCs (from Figure 1, without transplantation). Note that genes highly expressed in pre-graft TA MuSCs (from Figure 1, coloured in pink) were mostly upregulated in EOM MuSCs following grafting, while genes highly expressed in pre-graft EOM MuSCs (from Figure 1, coloured in blue) were mostly downregulated in EOM MuSCs following grafting. (C) PCA analysis of TA pre-graft, EOM pre-graft and EOM post-graft MuSCs using the expression values of genes differentially expressed between EOM and TA pre-graft MuSCs. PC1 separates EOM and TA MuSCs and shows that after engrafting EOM MuSCs into the TA muscle, they resemble transcriptionally TA MuSCs more than EOM MuSCs. (D) (left) Classification of EOM and TA-specific genes as resistant, responsive, or intermediate upon heterotopic transplantation of EOM MuSCs. Determining the responsiveness of each gene to transplantation was carried out in two steps. First, expression values for TA-specific genes (resp EOM-specific) in pre-graft MuSCs were rescaled to 0-100, where 0 represented the mean expression in EOM (resp TA) MuSCs and 100 represented the mean expression in TA (resp EOM) MuSCs. Next, the expression of EOM and TA-specific genes were rescaled similarly in post-graft EOM MuSCs. Each dark blue dot represents a gene with corresponding rescaled value in post-graft EOM MuSCs. EOM-specific genes with rescaled values in post-graft EOM MuSCs less than 25 were classified as responsive, between 25 and 75 as intermediate and above 75 as resistant. TA-specific genes with rescaled values in post-graft EOM MuSCs less than 25 were classified as resistant, between 25 and 75 as intermediate and above 75 as responsive. (right) Distribution of responses of EOM and TA-specific genes in post-graft EOM MuSCs. (E) Expression of selected markers between TA pre-graft, EOM pre-graft and EOM post-graft MuSCs. Many TA marker genes were upregulated to TA-like levels when EOM MuSCs were grafted into TA muscle. In addition, some EOM marker genes such as *Lmx1a* and *Mobp* were downregulated in this scenario. *** p < 0.001, ** p < 0.01, * p < 0.05 by Welch’s t test. (F) Gene expression analysis throughout the *HoxA, HoxB, HoxC* and *HoxD* clusters in post-graft EOM (left) and TA (middle) MuSCs. Genes were color-coded according to their antero-posterior expression domain in mouse at embryonic day 12.5 (right, adapted from^57^). All TA-specific *Hox* genes were upregulated in post-graft EOM MuSCs.

To evaluate exclusively the response of EOM MuSCs to a heterologous microenvironment while setting aside modifications in transcriptome due to the transplantation process, we calculated a correction coefficient based on post-graft TA samples and applied it to the transcriptome of post-graft EOM and post-graft TA samples (see Methods). As expected, the corrected post-graft TA MuSCs clustered with pre-graft TA MuSCs (Supplementary Figure 3E). Post-graft EOM MuSCs were then corrected similarly and hereafter compared to pre-graft EOM and TA MuSCs (Figures 3B to 3F). We observed a global shift in the transcriptome of post-graft EOM MuSCs towards a TA profile (Figure 3B). The majority of TA-specific genes were upregulated in post-graft EOM MuSCs while EOM-specific genes were downregulated (Figure 3B), indicating that a large proportion of the transcriptome was remodeled by the cellular microenvironment. To further investigate these changes, we performed PCA analysis using location-specific DEGs and observed that post-graft EOM MuSCs tended to cluster closer to pre-graft TA MuSCs than to pre-graft EOM MuSCs (PC1, Figure 3C). The higher similarity of the post-graft EOM samples to pre-graft TA MuSCs was further corroborated by transcriptomic correlation and hierarchical clustering analysis (Supplementary Figure 3F). We then classified location-specific genes in post-graft EOM MuSCs according to their degree of change into three categories: responsive, intermediate, and resistant (Figure 3D left). Genes were classified as resistant if they maintained their initial expression level in EOM MuSCs post-graft, while genes were classified as responsive if they adopted a TA-like expression level in EOM MuSCs post-graft. More specifically, responsive genes included EOM-specific genes fully downregulated to pre-graft TA level or TA-specific genes fully upregulated to pre-graft TA level; intermediate genes were those whose expression levels were between pre-graft EOM and TA samples; and resistant genes included EOM-specific genes remained at pre-graft EOM level or TA-specific genes remained at pre-graft EOM level. In agreement with the observed global shift of the transcriptome (Figure 3B), the majority of the location-specific genes showed a responsive or intermediate profile (∼90% of the EOM-specific DEGs and ∼80% of TA-specific DEGs) while only around 10-20% of DEGs were resistant after transplantation (Figure 3D right). Importantly, post-graft EOM MuSCs upregulated the TA-specific markers *Pax3, Tbx1 and Lbx1* while they down-regulated the EOM-specific markers *Alx4, Lmx1a and Mobp* (Figure 3E), confirming their acquisition of a limb-like transcriptional identity. Of note, the EOM-enriched *Pitx2* gene was resistant to transplantation (Figure 3E). Strikingly, all TA-specific *Hox* genes were upregulated in post-graft EOM MuSCs, with intermediate or responsive behaviours (Figure 3F). This observation is in agreement with previous reports showing that *Hox*-negative tissues or cells adopt the *Hox* status of their new location upon grafting^62,63^.

We then followed a similar approach to investigate the response of the epigenome to the different cellular microenvironments in the absence of confounding factors. We first calculated a correction coefficient based on the average DNA methylation differences between the methylomes of pre-graft and post-graft TA MuSCs using windows covering the promoter and enhancer regions and applied it to the methylome of the post-graft MuSCs. The correction coefficient was applied to promoters and enhancers of post-graft TA and EOM MuSCs. After correction, we performed PCA analysis considering promoters of location-specific genes (DEGs). Consistent with our previous results (Figure 2B), we observed no substructure of the data, suggesting no clear changes in DNA methylation at the promoter level (Figure 4A). However, a clear pattern was observed at enhancer regions associated with EOM and TA DEGs. PCA and clustering analysis of these regulatory regions showed that post-graft EOM MuSCs clustered closer to TA samples than to EOM samples (Figure 4B and Supplementary Figure 4A) indicating that enhancer regions undergo global reprogramming after transplantation in response to the cellular microenvironment. Overall, enhancers associated with EOM DEGs gained DNA methylation whereas those associated with TA DEGs lost DNA methylation in post-graft EOM MuSCs (Figure 4C). These epigenetic changes could contribute to some of the observed gene expression changes upon heterotopic transplantation of EOM MuSCs. Some examples include the EOM DEGs *Eya2* and *Rtn1*, which gain DNA methylation at their associated enhancer and become repressed, and the TA DEGs *Cit* and *Enpep* which lose DNA methylation at their associated enhancer and become more expressed in the post-graft EOM MuSCs (Figure 4D). Interestingly, the TA-specific genes *Lbx1* and *Pax3* show moderate changes of methylation at their associated enhancers while they become more expressed, indicating that additional mechanisms regulate changes of expression observed in EOM MuSCs upon heterologous transplantation. Of note, changes of enhancer methylation were also observed for some genes showing little variation in expression upon grafting, as for *Myod* and *Pitx2* (Supplementary Figure 4B). Interestingly, the *HoxA* gene cluster acquired a DNA methylation pattern resembling that of TA MuSCs (Figure 4E), associated with an overall increased expression level (Figure 3F). The other *Hox* clusters did not show much change in their DNA methylation patterns upon heterologous transplantation (Supplementary Figure 4C). This was not unexpected as EOM and TA MuSCs were found to have similar DNA methylation patterns across these regions. For the *HoxC* cluster, other mechanisms may be responsible for the expression difference observed between EOM and TA MuSCs (Figure 1E) and the gain of expression seen in EOM MuSCs after heterologous transplantation (Figure 3F). For the *HoxB* and *HoxD* clusters, the relatively low methylation levels across these regions may be responsible for the lack of expression in EOM MuSCs, TA MuSCs, and EOM MuSCs after heterologous transplantation (Figures 1E and 3F).

**Figure 4.**
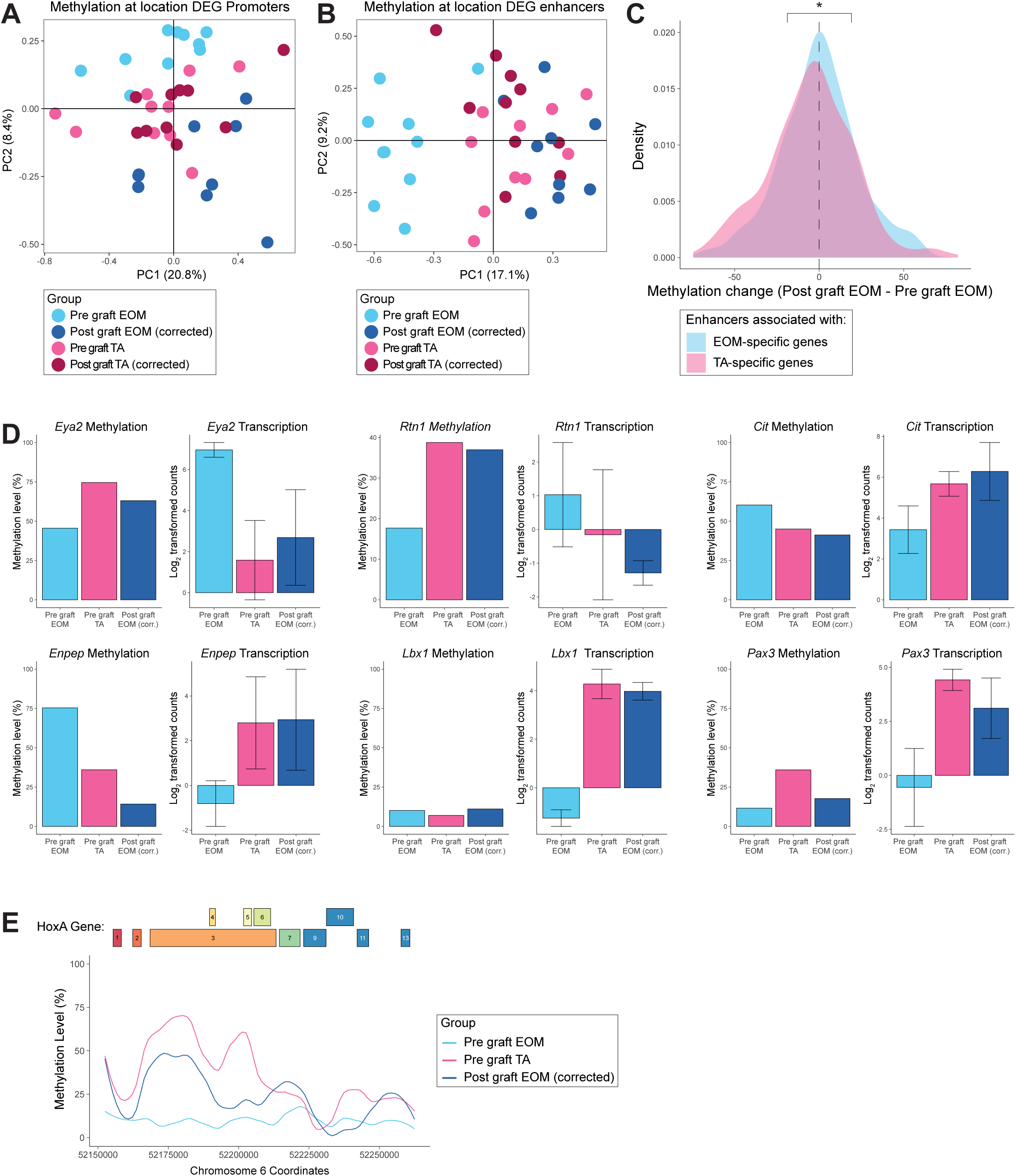
DNA methylation at enhancers partially accounts for the transcriptome plasticity of MuSCs upon heterotopic transplantation. (A) PCA analysis of TA pre-graft, EOM pre-graft and EOM post-graft MuSCs DNA methylation at promoters fails to separate samples based on anatomical location. (B) PCA analysis of TA pre-graft, EOM pre-graft and EOM post-graft MuSCs DNA methylation at enhancers separates samples based on anatomical location and demonstrates that EOM MuSCs after grafting resemble TA MuSCs at an epigenetic level in the enhancer context. (C) Density plots of the methylation difference between post-graft EOM MuSCs and pre-graft EOM MuSCs at enhancers associated with EOM or TA upregulated genes. EOM enhancers became hypermethylated in EOM MuSCs after grafting into the TA environment, while TA enhancers were hypomethylated. * p < 0.05 by Student’s t test. (D) Enhancer methylation and gene expression levels of selected EOM and TA DEGs that are responsive to grafting. (E) DNA methylation level across the *HoxA* gene cluster in pre-graft EOM, pre-graft TA and post-graft EOM samples. The *HoxA* region was highly methylated in TA MuSCs but not in EOM MuSCs. Notably, DNA methylation was gained across this region when EOM MuSCs were grafted into TA muscle.

## DISCUSSION

Here we examined the combined transcriptional and DNA methylation signatures of head (EOM) and limb (TA)-derived MuSCs to assess the molecular determinants associated with their diversity and inform on their functional differences such as susceptibility to disease and regenerative capacity. In this context, we also investigated the extent to which a head-specific identity was determined cell-autonomously through heterotopic transplantation into a limb environment, followed by transcriptome and DNA methylation profiling of re-isolated cells.

Consistent with previous reports, we identified known EOM (*e.g. Alx4, Pitx2*) and TA (*e.g. Pax3, Lbx1, Hox* genes)-specific genes. Surprisingly, we observed expression of *HoxA* and *HoxC* genes, while limb development has been reported to rely mostly on *HoxA* and *HoxD* clusters, with minor contributions from the *HoxB* and *HoxC* clusters^64^. However, we detected expression of *Hoxa11* but not *Hoxa13* in TA MuSCs, consistent with their differential roles in the proximal-to-distal patterning of the limb^64^. Interestingly, we identified *Tbx1* as a TA-specific gene, although it is required for specification of branchial-arch derived craniofacial muscles^30^. This observation might also result from the different fiber-type composition of EOM and TA muscles and the specific properties of myogenic progenitors derived from different muscle fiber-types^65^. Additionally, EOM and TA-specific genes were enriched for developmental processes, which might reflect the persistence of molecular differences acquired during embryogenesis through adulthood. When and to what extent the transcriptomes of head and limb muscle progenitors initially diverge during embryogenesis remains to be addressed.

While several studies investigated the DNA methylation patterns of specific loci (*Myod*^66^, *Myogenin*^67^, *Desmin*^68^, *Six1*^69^, *α-smooth muscle actin*^70^) or genome-wide^43–45,71–77^ during myogenic specification or myoblast differentiation *in vitro*, we report, to the best of our knowledge, the first genome-wide DNA methylation profiling of pure adult quiescent MuSCs. The adult quiescent MuSCs we analysed here have much lower global methylation levels (∼50%) than somatic cells (∼70%)^78^, notably muscle fibers (∼75%)^79^, and other adult stem cells such as intestinal stem cells (78%)^80^ and hematopoietic stem cells (84%)^81^. Whether such a low methylation level is required for the stemness and quiescence properties of adult MuSCs remains to be explored, as well as the dynamics of DNA methylation between MuSCs and mature myofibers. Our analysis identified some distinct DNA methylation profiles between EOM and TA MuSCs, notably at enhancers associated with location-specific genes. These results suggest that location-specific transcriptome signatures are determined by location-specific DNA methylation patterns. We are not aware of any previous studies reporting a spatial address of enhancer methylation. It is conceivable therefore that the enhancer epigenome not only registers developmental and tissue-specificity but also the location (along the anterior-posterior axis in this case). Previous reports analysed DNA methylation patterns of similar tissue samples across several anatomical locations^82,83^, where changes in cellular composition might be a confounding factor. We report here for the first time the co-analysis of DNA methylome and transcriptome of pure populations of tissue-specific adult stem cells between different anatomical locations.

Our analysis of EOM and TA MuSCs re-isolated after engrafting each population into injured TA muscle followed by regeneration revealed a global reprogramming of EOM transcriptome towards a TA MuSC transcriptome, indicating that the extracellular environment strongly determines the location-specific signatures we identified. This observation is in agreement with a previous report^84^, indicating that stem cell-derived myogenic progenitors remodel partially their molecular signature towards an adult quiescent MuSC transcriptome upon *in vivo* engraftment. Importantly, our data show that TA MuSCs re-isolated after engrafting and regeneration display important transcriptome differences with their pre-grafting TA MuSCs counterparts. This suggests that comparing engrafted cells re-isolated after regeneration to adult quiescent MuSCs^84^ could underestimate the extent of transcriptome plasticity driven by the *in vivo* environment.

Significantly, we identified different classes of EOM and TA-specific genes in EOM MuSCs depending on their behaviour following *in vivo* engraftment: resistant, responsive, and intermediate. Genes with intermediate responses could reflect *i)* the stable acquisition of new intermediary expression states due to buffering from initial identity, *ii)* the existence of transcriptional changes with slow kinetics towards full reprogramming or *iii)* the existence of different subpopulations of EOM MuSCs with variable reprogramming efficiency. The first two hypotheses are favored due to the long-term transcriptional changes associated with our experimental procedure and the existence of fully-responsive genes, indicating a complete reprogramming of such genes in each cell.

Transplantation of cranial-derived stem cells allowed us to assess positional information, given that EOM MuSCs lack expression of *Hox* genes. Interestingly, we observed that all TA-specific *Hox* genes were upregulated in EOM engrafted MuSCs, reflecting the acquisition of the *Hox* status of their new location, as observed previously after transplantation of *Hox*-negative cells in a *Hox*-positive domain^63,85^ and unlike myoblasts maintaining their axial identity in culture^86^, indicating that factors derived from the *in vivo* environment might determine the plasticity of the *Hox* status. Whether the initial *Hox*-free status of EOM MuSCs is responsible for their superior regenerative capacity over *Hox*-positive limb-derived MuSCs remains to be determined. The molecular mechanisms of the *Hox* expression and methylation plasticity in adult stem cells remain to be identified, and their manipulation could be of interest in the context of regenerative medicine^87^.

We speculate that the limited set of EOM resistant genes could contribute to a niche-independent and cell-intrinsic high engraftment potential of EOM MuSCs. Interestingly, these genes include *Pitx2* and genes involved in thyroid hormone signalling (*Dio2*, encoding the type-II iodothyronine deiodinase, a thyroid hormone activator, and the Tsh hormone receptor *Tshr*). Of note, *Pitx2* overexpression was shown to increase the regenerative capacity of MuSCs^88^ and *Pitx2;Pitx3* double mutant mice have impaired muscle regeneration upon injury^89^, while thyroid hormone signalling was reported to be critical for MuSC survival and muscle regeneration^90,91^. Therefore, EOM resistant genes could be of interest as deterministic candidate regulators of EOM MuSC identity. Such determinants could be exploited for improving cellular therapy of muscular dystrophies^39,92,93^ using the abundant limb MuSCs.

Finally, we identified plastic DNA methylation patterns in engrafted EOM MuSCs, notably at enhancers associated with location-specific genes and at the *HoxA* gene cluster. Importantly, EOM enhancers were annotated based on H3K27ac ChIP data from limb-derived MuSCs^58^, which might not fully represent actual enhancers in EOM MuSCs. These observations indicate that cell-extrinsic cues are relayed through specific modifications of DNA methylation patterns to establish a MuSC transcriptome profile matching the new location. Future studies will be required to identify key molecular determinants of this interplay between the niche, the epigenome, and the transcriptome.

In summary, we report the molecular profiling of two populations of adult MuSCs which were reported to exhibit distinct regenerative capacities and disease susceptibility. We identify differences in the methylation of enhancers and demonstrate plasticity upon exposure to a new microenvironment. Therefore, the molecular characterization and functional analysis of MuSCs from multiple muscles and physiopathological conditions will be highly informative for a more complete understanding of muscle stem cell heterogeneity and plasticity.

**Supplementary Figure 1.**
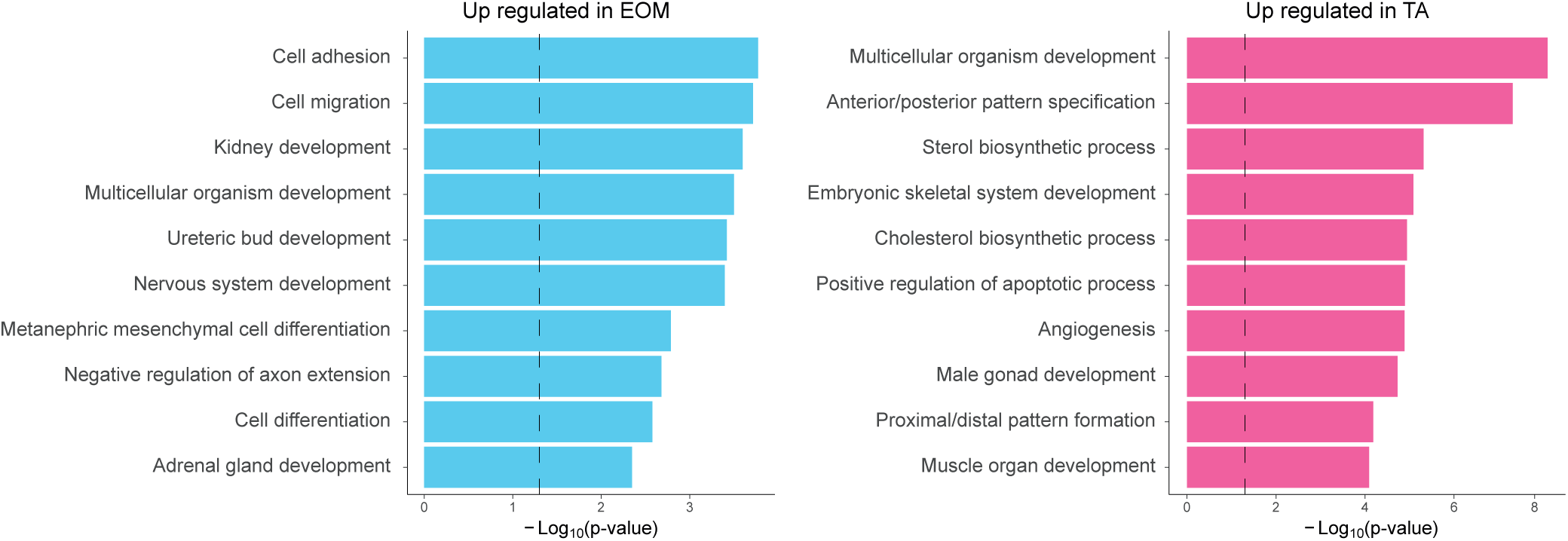
Gene ontology analysis of the genes upregulated in EOM or TA MuSCs. The 10 most significant categories are shown.

**Supplementary Figure 2.**
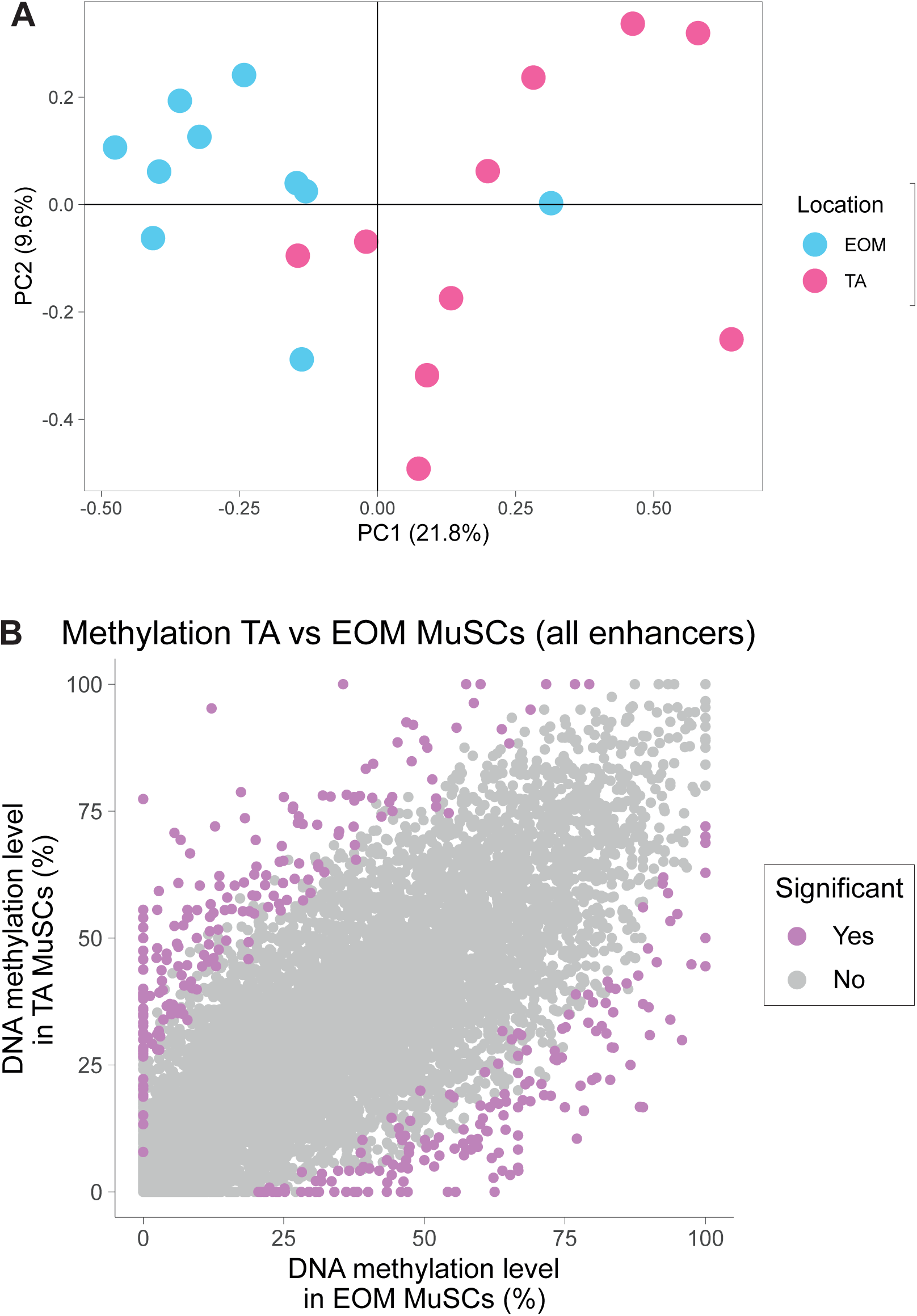
EOM and TA MuSCs display different DNA methylation patterns at enhancers associated with anatomical location-specific genes. (A) PCA analysis of DNA methylation of 544 enhancers associated with genes specifically expressed in TA or EOM MuSCs. EOM MuSCs and TA MuSCs separate according to their location. (B) Scatter plot comparing the mean DNA methylation levels of all enhancers in EOM and TA MuSCs. Enhancers were colour coded as significant if they were found to be differentially methylated by a rolling Z score approach (p < 0.05). See Figure 2E for genes with location-specific expression and enhancer methylation.

**Supplementary Figure 3.**
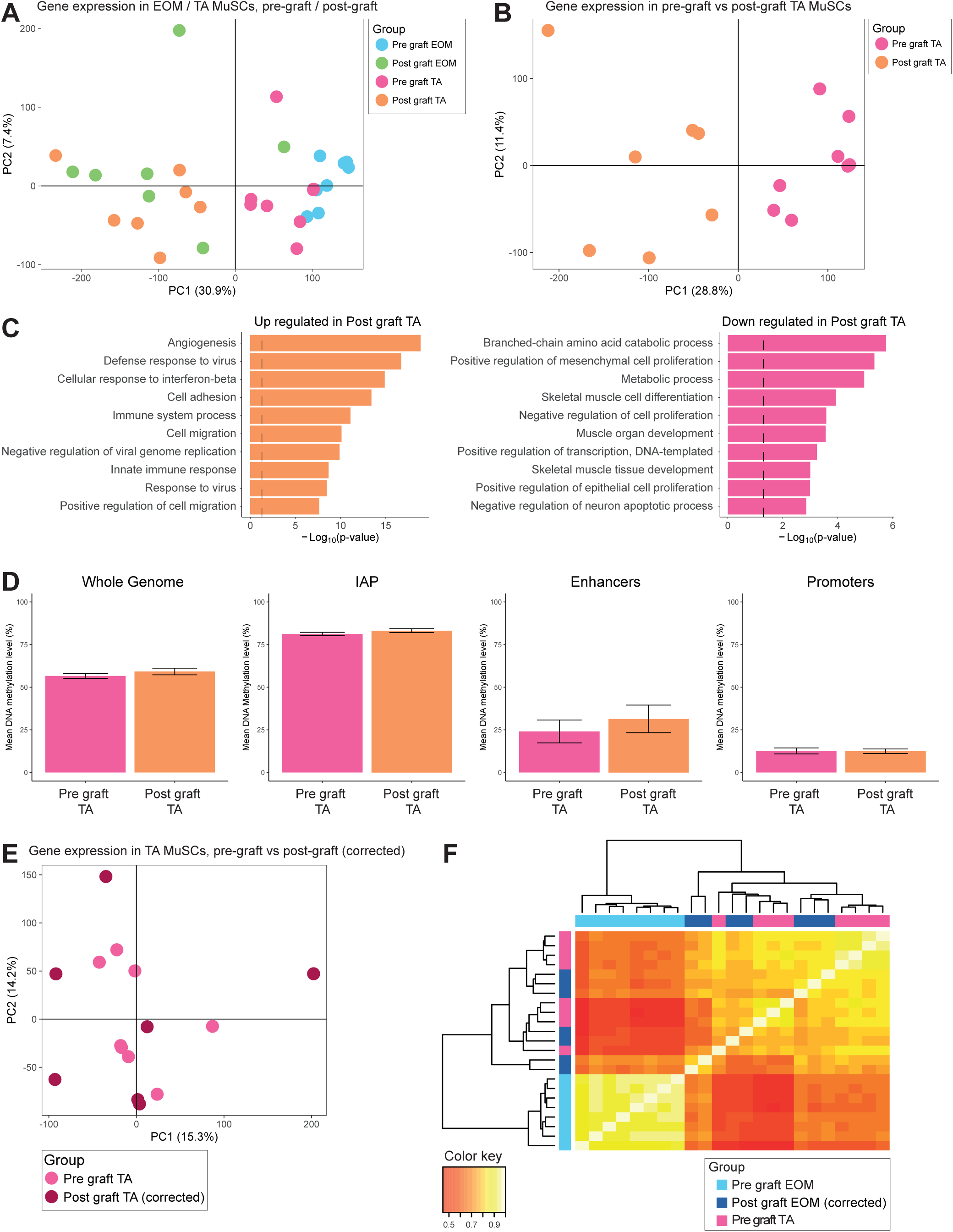
Transplantation of MuSCs induces long-term transcriptome and epigenome modifications. (A) Whole transcriptome PCA analysis of pre-graft and post-graft TA and EOM MuSCs. Samples separate into before and after grafting. (B) Whole transcriptome PCA analysis of pre-graft and post-graft TA MuSCs. Pre and post-graft samples cluster away from each other suggesting there is a residual impact from the grafting procedure even after the recovery period. (C) GO categories of genes upregulated (left) and downregulated (right) in post-graft *vs* pre-graft TA MuSCs. Upregulated categories suggest there was a remaining inflammatory response. (D) Mean DNA methylation levels in pre-graft and post-graft TA MuSCs for whole genome, repeat elements, promoters and enhancers. Overall there was a global increase in DNA methylation after grafting. (E) Whole transcriptome PCA analysis of pre-graft TA MuSCs and post-graft MuSCs after applying a correction coefficient accounting for transcriptome modifications specifically induced by the transplantation procedure (see Methods). Pre and post-graft samples no longer cluster separately. (F) Hierarchical clustering analysis using Euclidean distance of Pearson correlation values between TA pre-graft, EOM pre-graft and EOM post-graft MuSCs. The samples cluster separately based on anatomical location. Notably, after engrafting EOM MuSCs into TA muscle they cluster with TA MuSCs rather than EOM MuSCs.

**Supplementary Figure 4.**
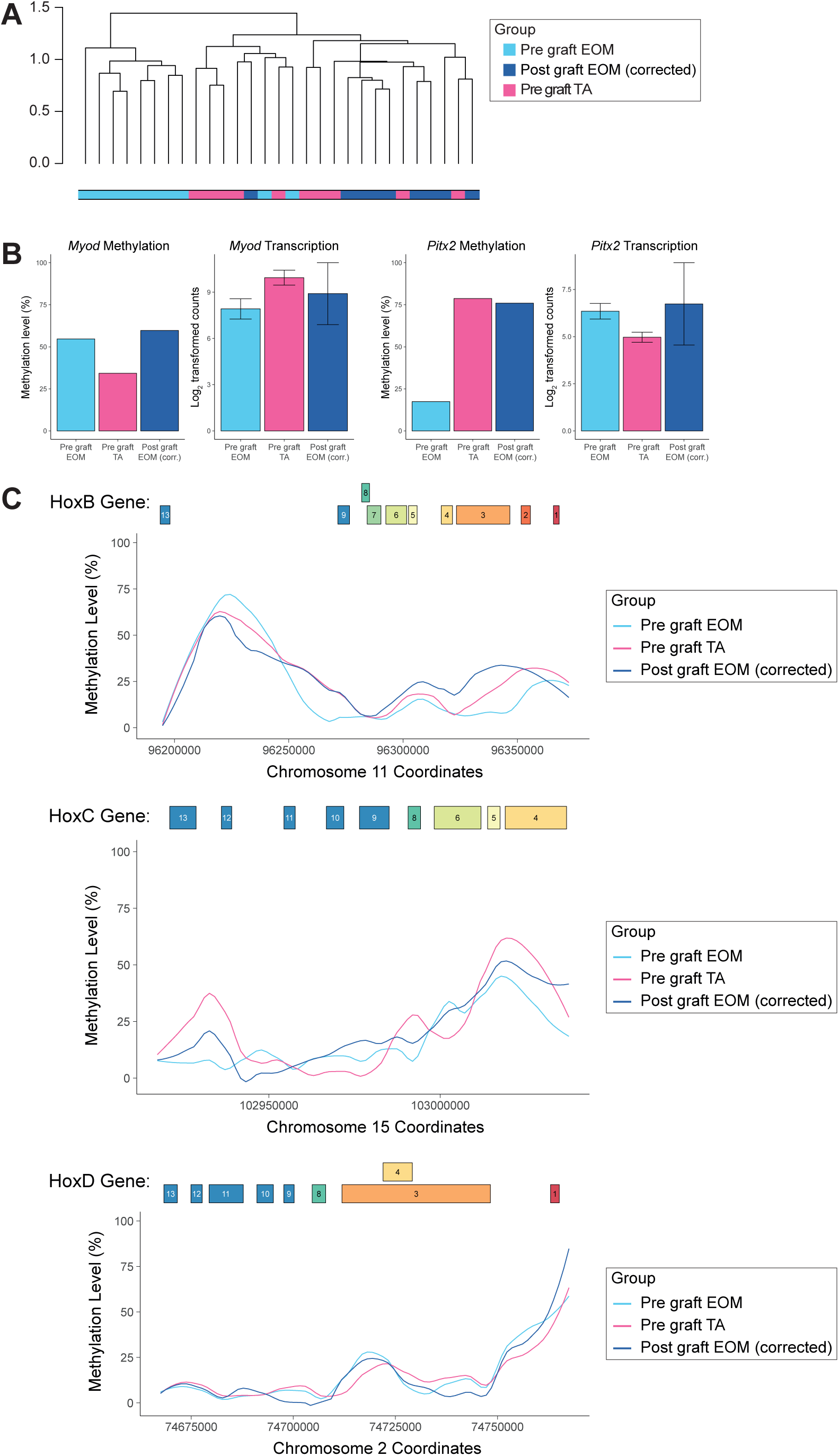
DNA methylation changes in MuSCs upon heterotopic transplantation. (A) Hierarchical clustering analysis using Euclidean distance of Pearson correlation values between TA pre-graft, EOM pre-graft and EOM post-graft MuSCs overall clusters samples based on location. Notably, post-graft EOM MuSCs cluster with TA MuSCs rather than with EOM MuSCs. (B) Enhancer methylation and gene expression levels of *Myod* and *Pitx2* prior to and following grafting. (C) DNA methylation level across the *HoxB* (top), *HoxC* (middle) and *HoxD* (bottom) gene clusters in pre-graft EOM, pre-graft TA and post-graft EOM samples. DNA methylation levels were similar between EOM MuSCs and TA MuSCs. In addition, grafting EOM MuSCs into the TA muscle environment did not substantially affect the DNA methylation across these regions.

**Supplementary Table 1.**
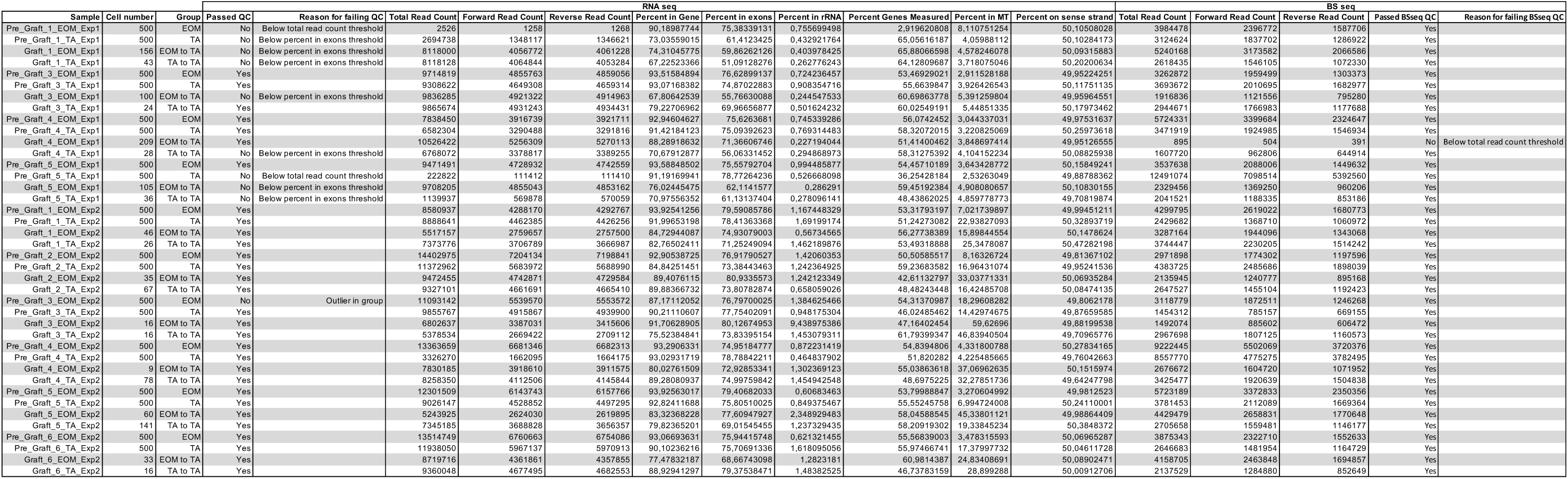
RNA-seq and BS-seq quality control.

## METHODS

### Mice

Animals were handled according to national and European Community guidelines and an ethics committee of the Institut Pasteur (CETEA) in France approved protocols. *Tg:Pax7-nGFP* ^17^ and *Rag2*^*-/-*^ *;γC*^*-/-* 59^ mice were used in this study, on C57BL/6;DBA2 F1/JRj and C57BL/6J genetic backgrounds respectively. 6 to 8-week-old male littermates were used.

### Isolation of muscle stem cells

Mice were sacrificed by cervical dislocation. *Tibialis anterior* and extraocular muscles were dissected and placed into cold DMEM (ThermoFisher, 31966). Muscles were then manually chopped with scissors and put into a 15 ml Falcon tube containing 10 ml of DMEM, 0.08% collagenase D (Sigma, 11 088 882 001), 0.1% trypsin (ThermoFisher, 15090), 10 µg/ml DNase I (Sigma, 11284932) at 37°C under gentle agitation for 25 min. Digests were allowed to stand for 5 min at room temperature and the supernatants were added to 5 ml of foetal bovine serum (FBS; Gibco) on ice. The digestion was repeated 3 times until complete digestion of the muscle. The supernatants were filtered through a 70-µm cell strainer (Miltenyi, 130-098-462). Cells were spun for 15 min at 515g at 4°C and the pellets were resuspended in 1 ml cold freezing medium (10% DMSO (Sigma, D2438) in FBS) for long term storage in liquid nitrogen or processed directly through FACS-isolation for transplantations.

Before isolation by FACS, samples were thawed in 50 ml of cold DMEM, spun for 15 min at 515g at 4°C. Pellets were resuspended in 300 µl of DMEM 2% FBS 1 µg/mL propidium iodide (Calbiochem, 537060) and filtered through a 40-µm cell strainer (BD Falcon, 352235). Viable muscle stem cells were isolated based on size, granulosity and GFP intensity using a MoFlo Astrios cell sorter (Beckmann Coulter).

Cells were collected in 5 µl cold RLT Plus buffer (Qiagen, 1053393) containing 1U/µl RNAse inhibitor (Ambion, AM2694), flash-frozen on dry ice and stored at −80°C, or in cold DMEM 2% FBS.

### Muscle stem cell transplantations

Muscle injury was done as described previously^94^. Briefly, mice were anesthetized with 0.5% Imalgene/2% Rompun. Both TAs of recipient immunocompromised *Rag2*^*-/-*^*;γC*^*-/-*^ mice were injured with 50 µl of 10 µM cardiotoxin (Latoxan, L8102) in NaCl 0.9% 24 h before transplantation. Muscle stem cells from freshly dissociated EOM and TA muscles of *Tg:Pax7-nGFP* mice were isolated by FACS, spun for 15 min at 515g at 4°C, resuspended in 10 µl of cold PBS and injected into recipient TAs. EOM and TA muscle stem cells from a donor mouse were injected into separate TAs of the same recipient mouse, and equivalent numbers of EOM and TA cells were injected. Transplanted muscle stem cells were re-isolated by FACS based on GFP positivity after three to four weeks.

### Bisulfite-seq and RNA-seq Library Preparation

RNA was separated from cell lysates and processed into libraries using the G&T method as described in^47^. DNA was also processed into libraries using the bulk protocol described in^48^. Bisulfite-seq libraries were sequenced on an Illumina HiSeq-2500 with 125bp paired end reads and RNA-seq libraries were sequenced on an Illumina HiSeq-2500 with 75bp paired end reads.

### Bisulfite-seq analysis

Adapter sequences and poor-quality calls were removed from the raw sequencing files using Trim galore (version 0.4.2). Reads were then mapped to mouse genome GRCm38 and deduplicated using Bismark (version 0.16.3).

Promoters were defined as −2000bp to 500bp of the TSS of Ensembl genes. Promoters were split into CpG Islands (CGI) promoters and non-CGI promoters based on whether they overlapped a CpG island. H3K27ac peaks were called using macs2 on H3K27ac ChIP-seq data obtained from^58^. Enhancers were defined as H3K27ac peaks that did not overlap promoters. Enhancers were linked to genes based on proximity. At least two CpG sites needed to be covered for a promoter or enhancer to be taken forward. In addition, promoters and enhancers needed to be covered in at least 3 samples of each group to be considered in the analysis.

### RNA-seq analysis

Adapter sequences and poor-quality calls were removed from the raw sequencing files using Trim galore (version 0.4.4). Reads were then mapped to mouse genome GRCm38 using Hisat2 (version 2.1.0). Log2 transformed counts were generated using Seqmonk. Differential expression analysis was carried out in R using DESeq and genes that demonstrated a fold change greater than 2 and a p-value less than 0.05 were classified as differentially expressed.

Correcting for the effect of transplantation was performed by initially determining the expression change for all genes between TA MuSCs before and after transplantation. This expression change was then deducted from all post-graft samples to generate corrected values that excluded the effect of transplantation alone.

## ACKNOWLEDGMENTS

We would like to thank the Flow Cytometry Platform of the Center for Technological Resources and Research (Institut Pasteur) and the Wellcome Trust Sanger Institute sequencing facility for assistance with Illumina sequencing. This project was supported by grants from Institut Pasteur, Agence Nationale de la Recherche (Laboratoire d’Excellence Revive, Investissement d’Avenir; ANR-10-LABX-73), Association Française contre les Myopathies (21857), CNRS, the European Research Council (Advanced Research Grant 332893), and the Biotechnology and Biological Sciences Research Council (BBSRC, CBBS/E/B/000C0425). I.H.-H. was supported by a Marie Sklodowska-Curie Individual Fellowship (751439).

## AUTHOR CONTRIBUTIONS

B.E., T.S., S.T. and W.R. proposed the concept and designed the experiments. B.E. and G.C. performed mouse dissections. B.E. and P.H.C. performed FACS. B.E. performed grafting experiments. T.S. performed library preparation and sequencing. D.G., I.H.H. and T.S. developed the analysis methodologies and analysed the experiments. B.E., D.G., I.H.H, S.T. and W.R. wrote the paper. All authors read and agreed on the manuscript.

## REFERENCES

1. von Maltzahn, J., Jones, A. E., Parks, R. J. & Rudnicki, M. A. Pax7 is critical for the normal function of satellite cells in adult skeletal muscle. Proc. Natl. Acad. Sci. 110, 16474–16479 (2013).

2. Sambasivan, R. et al. Pax7-expressing satellite cells are indispensable for adult skeletal muscle regeneration. Development 138, 3647–3656 (2011).

3. Lepper, C., Partridge, T. A. & Fan, C.-M. An absolute requirement for Pax7-positive satellite cells in acute injury-induced skeletal muscle regeneration. Development 138, 3639–3646 (2011).

4. Webster, C. & Blau, H. M. Accelerated age-related decline in replicative life-span of Duchenne muscular dystrophy myoblasts: implications for cell and gene therapy. Somat. Cell Mol. Genet. 16, 557–65 (1990).

5. Blau, H. M., Webster, C. & Pavlath, G. K. Defective myoblasts identified in Duchenne muscular dystrophy. Proc. Natl. Acad. Sci. U. S. A. 80, 4856–60 (1983).

6. Chakkalakal, J. V, Jones, K. M., Basson, M. A. & Brack, A. S. The aged niche disrupts muscle stem cell quiescence. Nature 490, 355–360 (2012).

7. Bernet, J. D. & Olwin, B. B. P38 MAPK signaling underlies a cell autonomous loss of stem cell self-renewal in aged skeletal muscle. 2, 265–271 (2015).

8. Sousa-Victor, P. et al. Geriatric muscle stem cells switch reversible quiescence into senescence. Nature 506, 316–321 (2014).

9. Cosgrove, B. D. et al. Rejuvenation of the aged muscle stem cell population restores strength to injured aged muscles Benjamin. Nat. Med. 20, 255–264 (2014).

10. Price, F. D. et al. Inhibition of JAK/STAT signaling stimulates adult satellite cell function. 20, 1174–1181 (2015).

11. Tierney, M. T. & Sacco, A. STAT3 signaling controls satellite cell expansion and skeletal muscle repair. 76, 211–220 (2012).

12. Evans, W. J. & Campbell, W. W. Sarcopenia and age-related changes in body composition and functional capacity. J. Nutr. 123, 465–8 (1993).

13. Brack, A. S. & Muñoz-Cánoves, P. The ins and outs of muscle stem cell aging. Skelet. Muscle 6, 1–9 (2016).

14. Jang, Y. C., Sinha, M., Cerletti, M., Dall’Osso, C. & Wagers, A. J. Skeletal Muscle Stem Cells: Effects of Aging and Metabolism on Muscle Regenerative Function. Cold Spring Harb. Symp. Quant. Biol. 76, 101–111 (2011).

15. Renault, V. et al. Regenerative potential of human skeletal muscle during aging. Aging Cell 1, 132–9 (2002).

16. Montarras, D. et al. Direct Isolation of Satellite Cells for Skeletal Muscle Regeneration. Science (80-.). 309, 2064–2067 (2005).

17. Sambasivan, R. et al. Distinct regulatory cascades govern extraocular and pharyngeal arch muscle progenitor cell fates. Dev. Cell 16, 810–821 (2009).

18. Tajbakhsh, S., Rocancourt, D., Cossu, G. & Buckingham, M. Redefining the genetic hierarchies controlling skeletal myogenesis: Pax-3 and Myf-5 act upstream of MyoD. Cell 89, 127–38 (1997).

19. Relaix, F., Rocancourt, D., Mansouri, A. & Buckingham, M. A Pax3/Pax7-dependent population of skeletal muscle progenitor cells. Nature 435, 948–953 (2005).

20. Kassar-Duchossoy, L. et al. Pax3/Pax7 mark a novel population of primitive myogenic cells during development. Genes Dev. 19, 1426–1431 (2005).

21. Noden, D. M. & Francis-West, P. The differentiation and morphogenesis of craniofacial muscles. Dev. Dyn. 235, 1194–1218 (2006).

22. Couly, G. F., Coltey, P. M. & Le Douarin, N. M. The developmental fate of the cephalic mesoderm in quail-chick chimeras. Development 114, 1–15 (1992).

23. Diogo, R. et al. A new heart for a new head in vertebrate cardiopharyngeal evolution. Nature 520, 466–473 (2015).

24. Kelly, R. G., Jerome-Majewska, L. A. & Papaioannou, V. E. The del22q11.2 candidate gene Tbx1 regulates branchiomeric myogenesis. Hum. Mol. Genet. 13, 2829–2840 (2004).

25. Nathan, E. et al. The contribution of Islet1-expressing splanchnic mesoderm cells to distinct branchiomeric muscles reveals significant heterogeneity in head muscle development. Development 135, 647–657 (2008).

26. Saga, Y. et al. MesP1: a novel basic helix-loop-helix protein expressed in the nascent mesodermal cells during mouse gastrulation. Development 122, 2769–78 (1996).

27. Gage, P. J., Suh, H. & Camper, S. A. Dosage requirement of Pitx2 for development of multiple organs. Development 126, 4643–4651 (1999).

28. Harel, I. et al. Distinct Origins and Genetic Programs of Head Muscle Satellite Cells. Dev. Cell 16, 822–832 (2009).

29. Gage, P. J., Rhoades, W., Prucka, S. K. & Hjalt, T. Fate maps of neural crest and mesoderm in the mammalian eye. Investig. Ophthalmol. Vis. Sci. 46, 4200–4208 (2005).

30. Comai, G. & Tajbakhsh, S. Molecular and Cellular Regulation of Skeletal Myogenesis. in Current topics in developmental biology 110, 1–73 (2014).

31. Zammit, P. S. et al. Muscle satellite cells adopt divergent fates. J. Cell Biol. 166, 347–357 (2004).

32. Olguin, H. C. & Olwin, B. B. Pax-7 up-regulation inhibits myogenesis and cell cycle progression in satellite cells: a potential mechanism for self-renewal. Dev. Biol. 275, 375–88 (2004).

33. Conboy, I. M. & Rando, T. A. The regulation of Notch signaling controls satellite cell activation and cell fate determination in postnatal myogenesis. Dev. Cell 3, 397–409 (2002).

34. Brack, A. S. & Rando, T. A. Tissue-specific stem cells: lessons from the skeletal muscle satellite cell. Cell Stem Cell 10, 504–514 (2012).

35. Kaminski, H. J., Al-Hakim, M., Leigh, R. J., Bashar, M. K. & Ruff, R. L. Extraocular muscles are spared in advanced duchenne dystrophy. Ann. Neurol. 32, 586–588 (1992).

36. Schoser, B. G. H. & Pongratz, D. Extraocular Mitochondrial Myopathies and their Differential Diagnoses. Strabismus 14, 107–113 (2006).

37. Porter, J. D. et al. The sparing of extraocular muscle in dystrophinopathy is lost in mice lacking utrophin and dystrophin. J. Cell Sci. 111 (Pt 13), 1801–11 (1998).

38. Emery, A. E. The muscular dystrophies. Lancet 359, 687–695 (2002).

39. Stuelsatz, P. et al. Extraocular muscle satellite cells are high performance myo-engines retaining efficient regenerative capacity in dystrophin deficiency. Dev. Biol. 397, 31–44 (2015).

40. Ryall, J. G. et al. The NAD+-dependent sirt1 deacetylase translates a metabolic switch into regulatory epigenetics in skeletal muscle stem cells. Cell Stem Cell 16, 171–183 (2015).

41. Pallafacchina, G. et al. An adult tissue-specific stem cell in its niche: A gene profiling analysis of in vivo quiescent and activated muscle satellite cells. Stem Cell Res. 4, 77–91 (2010).

42. Liu, L. et al. Chromatin modifications as determinants of muscle stem cell quiescence and chronological aging. Cell Rep. 4, 189–204 (2013).

43. Tsumagari, K. et al. Early de novo DNA methylation and prolonged demethylation in the muscle lineage. Epigenetics 8, 317–332 (2013).

44. Carrió, E. et al. Deconstruction of DNA methylation patterns during myogenesis reveals specific epigenetic events in the establishment of the skeletal muscle lineage. Stem Cells 33, 2025–2036 (2015).

45. Miyata, K. et al. DNA methylation analysis of human myoblasts during in vitro myogenic differentiation: De novo methylation of promoters of muscle-related genes and its involvement in transcriptional down-regulation. Hum. Mol. Genet. 24, 410–423 (2015).

46. Hernando-Herraez, I. et al. Ageing affects DNA methylation drift and transcriptional cell-to-cell variability in mouse muscle stem cells. Nat. Commun. 10, 1–11 (2019).

47. Macaulay, I. C. et al. Separation and parallel sequencing of the genomes and transcriptomes of single cells using G&T-seq. Nat. Protoc. 11, 2081–2103 (2016).

48. Clark, S. J. et al. Genome-wide base-resolution mapping of DNA methylation in single cells using single-cell bisulfite sequencing (scBS-seq). Nat. Protoc. 12, 534–547 (2017).

49. Relaix, F. et al. Pax3 and Pax7 have distinct and overlapping functions in adult muscle progenitor cells. J. Cell Biol. 172, 91–102 (2006).

50. Gross, M. K. et al. Lbx1 is required for muscle precursor migration along a lateral pathway into the limb. Development 127, 413–424 (2000).

51. Brohmann, H., Jagla, K. & Birchmeier, C. The role of Lbx1 in migration of muscle precursor cells. Development 127, 437–445 (2000).

52. Kelly, R. G., Jerome-Majewska, L. A. & Papaioannou, V. E. The del22q11.2 candidate gene Tbx1 regulates branchiometric myogenesis. Hum. Mol. Genet. 13, 2829–2840 (2004).

53. Biressi, S. & Rando, T. A. Heterogeneity in the muscle satellite cell population. Seminars in Cell and Developmental Biology 21, 845–854 (2010).

54. Doucet-Beaupré, H., Ang, S. L. & Lévesque, M. Cell fate determination, neuronal maintenance and disease state: The emerging role of transcription factors Lmx1a and Lmx1b. FEBS Letters 589, 3727–3738 (2015).

55. Montague, P., McCallion, A. S., Davies, R. W. & Griffiths, I. R. Myelin-associated oligodendrocytic basic protein: A family of abundant CNS myelin proteins in search of a function. Developmental Neuroscience 28, 479–487 (2006).

56. Mallo, M., Wellik, D. M. & Deschamps, J. Hox genes and regional patterning of the vertebrate body plan. Developmental Biology 344, 7–15 (2010).

57. Pearson, J. C., Lemons, D. & McGinnis, W. Modulating Hox gene functions during animal body patterning. Nature Reviews Genetics 6, 893–904 (2005).

58. Machado, L. et al. In situ fixation redefines quiescence and early activation of skeletal muscle stem cells. Cell Rep. 21, 1982–1993 (2017).

59. Mazurier, F. et al. A Novel Immunodeficient Mouse Model-RAG2 gamma Cytokine Receptor Chain Double Mutants-Requiring Exogenous Cytokine Administration for Human Hematopoietic Stem Cell Engraftment Common. J. Interf. Cytokine Res. 19, 533–541 (1999).

60. Hardy, D. et al. Comparative Study of Injury Models for Studying Muscle Regeneration in Mice. PLoS One 11, e0147198 (2016).

61. De Micheli, A. J. et al. Single-Cell Analysis of the Muscle Stem Cell Hierarchy Identifies Heterotypic Communication Signals Involved in Skeletal Muscle Regeneration. Cell Rep. 30, 3583-3595.e5 (2020).

62. Itasaki, N., Sharpe, J., Morrison, A. & Krumlauf, R. Reprogramming Hox expression in the vertebrate hindbrain: influence of paraxial mesoderm and rhombomere transposition. Neuron 16, 487–500 (1996).

63. Leucht, P. et al. Embryonic origin and Hox status determine progenitor cell fate during adult bone regeneration. Development 135, 2845–2854 (2008).

64. Zakany, J. & Duboule, D. The role of Hox genes during vertebrate limb development. Current Opinion in Genetics and Development 17, 359–366 (2007).

65. Motohashi, N. et al. Tbx1 regulates inherited metabolic and myogenic abilities of progenitor cells derived from slow- and fast-type muscle. Cell Death Differ. 26, 1024–1036 (2019).

66. Brunk, B. P., Goldhamer, D. J. & Emerson, C. P. Regulated Demethylation of the myoD Distal Enhancer during Skeletal Myogenesis. 503, 490–503 (1996).

67. Lucarelli, M., Fuso, A., Strom, R. & Scarpa, S. The dynamics of myogenin site-specific demethylation is strongly correlated with its expression and with muscle differentiation. J. Biol. Chem. 276, 7500–7506 (2001).

68. Allen, M., Koch, C. M., Clelland, G. K., Dunham, I. & Antoniou, M. DNA methylation-histone modification relationships across the desmin locus in human primary cells. BMC Mol. Biol. 10, (2009).

69. Wu, W. et al. Core promoter analysis of porcine Six1 gene and its regulation of the promoter activity by CpG methylation. Gene 529, 238–244 (2013).

70. Hu, B., Gharaee-Kermani, M., Wu, Z. & Phan, S. H. Epigenetic regulation of myofibroblast differentiation by DNA methylation. Am. J. Pathol. 177, 21–28 (2010).

71. Illingworth, R. et al. A novel CpG island set identifies tissue-specific methylation at developmental gene loci. PLoS Biol. 6, e22 (2008).

72. Sørensen, A. L. et al. Lineage-specific promoter DNA methylation patterns segregate adult progenitor cell types. Stem Cells Dev. 19, 1257–1266 (2010).

73. Fernandez, A. F. et al. A DNA methylation fingerprint of 1628 human samples. Genome Res. 22, 407–419 (2012).

74. Calvanese, V. et al. A promoter DNA demethylation landscape of human hematopoietic differentiation. Nucleic Acids Res. 40, 116–131 (2012).

75. Nazor, K. L. et al. Recurrent Variations in DNA Methylation in Human Pluripotent Stem Cells and Their Differentiated Derivatives. Cell Stem Cell 10, 620–634 (2012).

76. Carrió, E. et al. Muscle cell identity requires Pax7-mediated lineage-specific DNA demethylation. BMC Biol. 14, 30 (2016).

77. Bigot, A. et al. Age-associated methylation suppresses SPRY1, leading to a failure of re-quiescence and loss of the reserve stem cell pool in elderly muscle. Cell Rep. 13, 1172–1182 (2015).

78. Lister, R. et al. Human DNA methylomes at base resolution show widespread epigenomic differences. Nature 462, 315–322 (2009).

79. Begue, G., Raue, U., Jemiolo, B. & Trappe, S. DNA methylation assessment from human slow- and fast-twitch skeletal muscle fibers. J. Appl. Physiol. 122, 952–967 (2017).

80. Kaaij, L. T. J. et al. DNA methylation dynamics during intestinal stem cell differentiation reveals enhancers driving gene expression in the villus. Genome Biol. 14, (2013).

81. Challen, G. A. et al. Dnmt3a and Dnmt3b have overlapping and distinct functions in hematopoietic stem cells. Cell Stem Cell 15, 350–364 (2014).

82. Kaz, A. M. et al. Patterns of DNA methylation in the normal colon vary by anatomical location, gender, and age. Epigenetics 9, 492–502 (2014).

83. Wu, M. et al. DNA methylation profile of psoriatic skins from different body locations. Epigenomics 11, 1613–1625 (2019).

84. Incitti, T. et al. Pluripotent stem cell-derived myogenic progenitors remodel their molecular signature upon in vivo engraftment. Proc. Natl. Acad. Sci. U. S. A. 116, 4346–4351 (2019).

85. Grapin-Botton, A., Bonnin, M. A., McNaughton, L. A., Krumlauf, R. & Le Douarin, N. M. Plasticity of transposed rhombomeres: Hox gene induction is correlated with phenotypic modifications. Development 121, 2707–21 (1995).

86. Grieshammer, U., Sassoon, D. & Rosenthal, N. A transgene target for positional regulators marks early rostrocaudal specification of myogenic lineages. Cell 69, 79–93 (1992).

87. Wang, K. C., Helms, J. A. & Chang, H. Y. Regeneration, repair and remembering identity: the three Rs of Hox gene expression. Trends in Cell Biology 19, 268–275 (2009).

88. Vallejo, D. et al. PITX2 Enhances the Regenerative Potential of Dystrophic Skeletal Muscle Stem Cells. Stem Cell Reports 10, 1398–1411 (2018).

89. L’Honoré, A. et al. The role of Pitx2 and Pitx3 in muscle 1 stem cells gives new insights into P38α MAP kinase and redox regulation of muscle regeneration. Elife 7, (2018).

90. Dentice, M. et al. The FoxO3/type 2 deiodinase pathway is required for normal mouse myogenesis and muscle regeneration. J. Clin. Invest. 120, 4021–4030 (2010).

91. Dentice, M. et al. Intracellular inactivation of thyroid hormone is a survival mechanism for muscle stem cell proliferation and lineage progression. Cell Metab. 20, 1038–1048 (2014).

92. McLoon, L., Thorstenson, K., Solomon, A. & Lewis, M. Myogenic precursor cells in craniofacial muscles. Oral Dis. 13, 134–140 (2007).

93. Kallestad, K. M. et al. Sparing of extraocular muscle in aging and muscular dystrophies: A myogenic precursor cell hypothesis☆. Exp. Cell Res. 317, 873–885 (2011).

94. Gayraud-Morel, B. et al. A role for the myogenic determination gene Myf5 in adult regenerative myogenesis. Dev. Biol. 312, 13–28 (2007).

